# GlycanPUG: WURCS-Based Heterogeneous Graph Pretraining for Structurally Ambiguous Glycans

**DOI:** 10.64898/2026.11.22.767969

**Authors:** Ye-fei Wang, Xiao-peng He, Wen-hao Zhang

**Affiliations:** East China University Of Science And Technology; Shanghai Jiao Tong University; East China University of Science and Technology

## Abstract

**Background:** Glycans orchestrate diverse biological processes, yet they remain underrepresented in computational biology compared to proteins and nucleic acids. Although WURCS 2.0 provides a machine-readable structural representation that preserves uncertainty, the evaluated IUPAC-based deterministic workflow processes 43.1 % of the GlyCosmos-derived cohort examined here, motivating approaches that retain structural ambiguity.

**Methods:** To overcome this limitation, GlycanPUG (Probabilistic Uncertainty-aware Graph) was introduced as a pretrained graph model for glycan representation learning. By leveraging WURCS 2.0, GlycanPUG represents candidate linkages and substituent attachments using weighted edges and uncertainty attributes within a heterogeneous graph framework. A 4-layer Heterogeneous Graph Neural Network with a dedicated node pre-enhancement module aggregates edge features weighted by the assigned edge weights, enabling representation of supported structurally ambiguous glycans without selecting a single resolved structure. The model is pretrained on a preprocessed subset of the GlyTouCan(Tiemeyer et al. 201 7) repository, which contained 253,596 entries before filtering. After excluding entries linked to downstream tasks and applying sequential structural filtering, 1 67,020 samples remained and were used for pretraining. Pretraining comprised nine self-supervised tasks.

**Results:** Benchmarking on the GlycanML dataset demonstrates that GlycanPUG shows competitive performance relative to GlycanGT across multi-label taxonomy classification and binary immunogenicity prediction. Crucially, on datasets natively containing structural uncertainties (GlyCosmos, GlyConnect), GlycanPUG effectively processes additional structurally ambiguous entries, achieving 83.5% processable coverage across all glycans from all biological sources in GlyCosmos, with taxonomic classification performance (e.g., mean macro F1 of 0.7376 for Domain). Ablation of the node pre-enhancement module reduces macro F1 , and post-hoc analysis identifies rank-dependent associations between structural uncertainty and prediction errors. All results reported above are averaged over ten independent random seeds (1 -1 0).

**Conclusions:** GlycanPUG broadens the set of glycans that can be represented while retaining supported structural ambiguities.

## 1 Introduction

Glycans orchestrate a vast array of biological processes, including immune regulation (Pinho *et al*. 2023, Tvaroška 2025, Rabinovich *et al*. 2026), viral entry mechanisms (Li *et al*. 2021 , Dugan, E., and Peiffer 2022, Spillings *et al*. 2022), cancer progression(Hu *et al*. 2023, Pinto and Parameswaran 2023, Pienkowski *etal.* 2025), and protein–glycan interactions (Carpenter *et al*. 2022, Lundstrøm *etal.* 2022, Canner, Schnaar, and Gray 2025); yet they remain underrepresented in computational biology compared to nucleic acids and proteins. This disparity stems not from a lack of biological relevance, but from the absence of a universally adopted, machine-readable structural representation that bridges glycobiology and bioinformatics (Bojar and Lisacek 2022).

The conventional IUPAC (International Union of Pure and Applied Chemistry) condensed notation, while human-readable, is inherently non-canonical. The same structure can have multiple valid strings, complicating string-based deduplication and comparison. WURCS 2.0 (Matsubara et al. 201 7) (Tanaka et al. 201 4) addresses these limitations by providing a canonical representation of the encoded structural information and natively supporting uncertain linkages and substituents. Although RoBERTa (Liu et al. 201 9)-based models pretrained on masked WURCS strings have been applied to glycan tasks with promising results (Shinkawa et al. 2022), sequence-based inputs did not explicitly encode graph connectivity. A heterogeneous graph representation was therefore adopted, retaining supported candidate connections and unresolved structural attributes. However, deterministic preprocessing may exclude entries with unresolved structural features, reducing coverage in the evaluated datasets.

To overcome this limitation, GlycanPUG (Probabilistic Uncertainty-aware Graph) was introduced as a pretrained graph model for glycan representation learning. The framework combines WURCS-derived candidate connections, direction-specific linkage features, and edge-weighted node pre-enhancement within a heterogeneous graph encoder. This representation supports pretraining on a filtered subset of the GlyTouCan (Tiemeyer et al. 201 7) repository. Benchmarking on the GlycanML (Xu, Geng *etal.* 2025) dataset demonstrates that GlycanPUG’s performance is competitive with that of the evaluated deterministic baseline across multi-label taxonomy classification and binary immunogenicity prediction, using the confidence-interval–based comparison protocol described in 2.7. On datasets natively containing structural uncertainties, additional ambiguous entries are processed. Ablation supports the contribution of the node pre-enhancement module.

## 2 Methods

### 2.1 Data

Data for pretraining and downstream evaluation were collected from several public repositories. Pretraining glycans were obtained from the GlyTouCan(Tiemeyer et al. 201 7) repository (accessed on 1 6 May 2026), comprising 253, 596 entries before preprocessing. The framework was evaluated on four downstream tasks: two glycan taxonomy classification tasks (using distinct datasets), one immunogenicity prediction task, and one linkage type classification task. For all tasks, pretrained GlycanPUG was used as a fixed feature extractor, and only task-specific classification heads were trained. For fair comparison with established baselines, the taxonomy dataset and the immunogenicity dataset were sourced from GlycanML(Xu, Geng et al. 2025). The second taxonomy dataset was sourced from GlyCosmos(Yamada et al. 2020). Data for linkage type classification (distinguishing N-linked, O-linked, and free oligosaccharides) were retrieved from GlyConnect(Alocci et al. 201 9).

### 2.2 Data Preprocessing

The preprocessing excluded the following categories from the pretraining corpus: samples overlapping with downstream tasks (removed by exact match on canonical WURCS 2.0 strings to prevent leakage); glycans with repeating structural units (indeterminate copy number, high heterogeneity); entries with uncertain notation in the Unit Count region—the field recording essential compositional unit information, whose ambiguity corrupts graph initialisation and feature counting (Appendix A.4); samples carrying rare linkages, dominated by nitrogen-mediated ones, which form atypical three-residue-on-one-nitrogen topologies requiring a dedicated nitrogen node type and bespoke edge definitions that would suffer severe class imbalance at such low abundance; samples with excessive uncertain connectivity, judged by the ratio of confirmed linkages (Appendix A.4) to total inter-residue glycosidic edges (Appendix C); and entries retaining rare features after iterative low-frequency token filtering across all feature categories (Appendix C), which repeatedly drops sub-threshold tokens to tighten the vocabulary. Per-category counts and thresholds are given in Appendix F.

For downstream data (see Section 2.1 ), preprocessing was unified across tasks to ensure consistency. To address class imbalance and ensure sufficient training signal, taxonomic labels or linkage types with fewer than 30 occurrences were filtered out prior to splitting. Regarding data partitioning, official splits from GlycanML were adopted for the first taxonomy and immunogenicity tasks to align with baselines. For the second taxonomy dataset, a multilabel stratified splitting strategy was implemented to preserve the overall label distribution across training, validation, and test sets, because structure-based clustering can concentrate rare labels in individual subsets; this choice preserves label balance but does not ensure separation of structurally similar glycans. All glycans were standardized to WURCS 2.0 (Matsubara et al. 201 7). Specifically, since glycans from GlycanML are natively represented in IUPAC notation, they were converted to WURCS 2.0 via the GlycanFormatConverter (Tsuchiya, Yamada, and Aoki-Kinoshita 201 9) (see Appendix B for detailed conversion logic). In contrast, native WURCS 2.0 structures from GlyConnect were directly retrieved using GlyTouCan IDs to preserve structural fidelity and avoid conversion errors.

Throughout downstream preprocessing, samples exhibiting anomalous linkages, non-standard repetitive units, unsupported uncertain notations, or conversion failures were excluded. Detailed experimental settings and the rationale for dataset selection are discussed in Section 3.2.

### 2.3 Graph Construction and Model Architecture

Due to the complexity of the WURCS 2.0 parsing (see Appendix A for syntax preliminaries) and feature extraction pipeline, the full graph construction procedure is detailed in Appendix C. In brief, supported alternative connections are expanded into candidate edges, while unresolved positions remain encoded as edge attributes. Each edge also carries an assigned scalar weight. This distinguishes candidate connectivity from uncertainty in the attachment position. For edge directionality, reverse edges of inter-residue glycosidic connections are explicitly constructed with orientation-specific types (e.g., “m-n” and “n-m”), whereas reverse edges for substituent attachments are automatically added due to their symmetric nature. The elements of the graph, their associated features, pre-processing steps prior to graph attention network (GAT) (Veličković et al. 201 8) input, and corresponding embedding dimensions are summarized in Fig. S.1 of Appendix C.

The encoder uses four heterogeneous graph attention layers with four attention heads per layer. Head outputs are combined by addition rather than concatenation. ReLU serves as the activation function, complemented by Dropout (rate=0.2) and Layer Normalization integrated with residual connections.

Prior work (Gong and Cheng 201 9, Chen and Chen 2021 ) incorporates edge information into graph representation learning. Building on this approach, a node pre-enhancement module precedes message passing in each heterogeneous graph neural network layer. The module aggregates incident-edge features using their assigned weights and fuses the aggregate with the corresponding node representation. This module incorporates incident-edge information before neighbor-node representations are combined during message passing.

### 2.4 Pretraining Setup

To learn glycan structural representations, nine masked attribute and edge-reconstruction tasks are used, spanning node-level (e.g., carbon number, anomeric configuration, substituent prediction) and edge-level (e.g., residue edge type) objectives. Detailed configurations—including mask ratios, loss functions, and task-specific masking logic such as co-masking strategies for cyclic detection—are comprehensively listed in Appendix D. For optimization, the model was trained using the AdamW optimizer (lr=1 ×1 0⁻³, weight decay=1 ×1 0⁻⁴). Learning rate scheduling was handled by ReduceLROnPlateau (patience=5, factor=0.5, min_lr=1 ×1 0⁻⁶), which reduces the rate when validation loss plateaus. Model convergence was determined via early stopping (patience=20 epochs), utilizing the 5-step EMA of the validation MLM loss (min delta=1 ×1 0⁻⁴) to mitigate noise and ensure stable termination.

### 2.5 Pooling and Graph Embedding

Pooling is performed separately for the two node types—Glycosyl Residue Node (representing sugar rings) and Substituent Node (representing substituents)—using PyTorch Geometric’s global mean pool and global max pool functions (Fey and Lenssen 201 9). With a configured hidden dimension of 256, each pooling operation outputs a 256-dimensional vector; concatenating the mean and max pooling results thus produces a 51 2-dimensional representation per node type. The pooled Glycosyl Residue Node (51 2D) and Substituent Node (51 2D) vectors are concatenated to form a 1 024-dimensional graph embedding, which is further appended with the 3-dimensional global features (UniqueRES count, RES count, and true linkage count) to yield the final 1 027-dimensional output. The LIN count tallies all linkage declarations in the WURCS linkage area, including uncertain substituent attachments whose host residue is undefined. The true linkage count is obtained by excluding from the LIN count those uncertain substituent attachments whose host residue is undefined, thereby retaining inter-residue linkage declarations, which may still contain positional or connectivity uncertainty. During feature encoding, any token not present in the predefined vocabulary is assigned to the designated unknown token. If no such token exists, it is mapped to the most frequent token in the vocabulary as a fallback encoding.

### 2.6 Downstream Task Adaptation

For all downstream tasks, pretrained models GlycanPUG and, where applicable, GlycanGT (Kitani et al. 2026) served as fixed feature extractors with frozen parameters; only task-specific classification heads were trained.

For the binary immunogenicity task on the GlycanML benchmark and the three-class glycosylation type task (N-linked, O-linked, free oligosaccharides) on GlyConnect, output layers produced 2D and 3D vectors, respectively, both with CrossEntropyLoss. The GlyConnect task and the GlyCosmos taxonomy task (Section 3. 4) were performed exclusively with GlycanPUG, because the retained WURCS inputs include structural ambiguities not supported by the evaluated deterministic preprocessing workflow.

For taxonomy and immunogenicity tasks on the GlycanML benchmark involving GlycanGT as a comparative baseline, a dual-branch architecture was adopted: the IUPAC branch utilizes GlycanGT (embedding dimension: 765), while the WURCS2.0 branch employs GlycanPUG (embedding dimension: 1 027). Both branches share subsequent hidden dimensions of 51 2 and 256 before mapping to class-specific output dimensions at each hierarchical level.

The downstream training configuration is described below. The multilabel taxonomy heads (GlycanML and GlyCosmos; eight ranks) used the Class-Balanced Focal configuration below; the binary immunogenicity and three-class glycosylation-type heads used standard CrossEntropyLoss with inverse-frequency class weights over softmax outputs. To mitigate class imbalance, a Class-Balanced variant of Focal Loss (Lin *et al*. 201 7) was used, with class-balanced weights (Cui *et al*. 201 9) derived from the effective number of samples and rescaled so that the most frequent class receives a weight of one, acting as the α factor on the positive term only (the negative term was left unweighted); the decay coefficient β was set to 0.9, 0.99 or 0.999 according to corpus size, and the focusing parameter was fixed at γ = 2.0. Training was monitored using EMA-smoothed validation macro-averaged AUC, with early stopping (patience: 68 epochs). A three-stage learning rate decay was implemented: if no significant improvement was observed for 1 8, 36, or 54 consecutive epochs, the learning rate was reduced by factors of 0.5, one-third, and one-quarter, respectively, with a minimum cap of 1 ×1 0⁻⁶. Threshold optimization was performed per class on the validation set via two-phase search: coarse scanning over [0.1 , 0.9] (step: 0.05), followed by fine-grained calibration within ±0.1 (step: 0.01 ). Inference used the resulting class-specific optimal thresholds. On the GlyCosmos dataset, this same configuration was retained to evaluate GlycanPUG on structurally ambiguous inputs.

### 2.7 Statistical Comparison

For each downstream task, model performance was evaluated over ten independent random seeds (1 –1 0). To compare any two models, the paired difference (candidate model − baseline model) was computed per seed, and a two-sided 95% confidence interval (CI) for the mean difference was constructed using the t-distribution. A three-way classification was applied: if the entire CI lay strictly above zero, the candidate model was classified as “better”; if the entire CI lay strictly below zero, the baseline model was classified as “better”; otherwise, the result was “inconclusive”. This protocol was uniformly applied across all comparative evaluations in this study.

### 2.8 Association Between Structural Uncertainty and Prediction Errors

To assess the association between structural uncertainty and prediction errors, a post-hoc analysis was conducted on the GlyCosmos test sets (reproduced via ten independent seeds; see Section 2. 7). Six graph-level uncertainty features were extracted from the parsed WURCS 2. 0 representation for each sample, corresponding to the six categories visualized in Fig. 6: inter-residue edge (uncertain residue connectivity), substituent attachment edge (uncertain substituent placement), ring bond type (ring connectivity uncertainty), residue edge type (linkage category uncertainty), anomeric (stereochemical uncertainty), and Carbon descriptor (carbon-level uncertainty). Detailed extraction logic is provided in Appendix E. A sample was counted as correctly classified only when its predicted multi-label vector matched the ground-truth vector exactly (zero Hamming distance across all labels at that rank); any single-label discrepancy was treated as a misclassification. Distributions were compared between correctly classified and misclassified samples using the one-sided Mann–Whitney U test. Effect sizes were quantified using Cohen’s d (absolute values plotted), and single-feature discrimination between correct and incorrect predictions was assessed using AUROC (with scores inverted for certainty-type metrics to maintain consistent directional interpretation). This analysis was restricted to the GlyCosmos taxonomy task; the glycosylation type classification task on GlyConnect was excluded because near-ceiling accuracy yielded too few misclassified samples for statistically meaningful comparison.

## 3 Results

### 3.1 Data

The downstream evaluation comprises four tasks: two multi-label taxonomy classification tasks using GlyCosmos (Yamada *et al*. 2020) and GlycanML (Xu, Geng *et al*. 2025), one immunogenicity binary classification, and one glycosylation type classification. As summarized in Fig. 2, the GlyCosmos taxonomy dataset contains 1 1 ,720–1 5,431 samples per hierarchical level (average ∼1 3,899) with 4–85 labels across 8 ranks, while the GlycanML taxonomy dataset includes 2,439–5,974 samples per level (average ∼4,958) with 3–64 labels. The immunogenicity and glycosylation tasks contain 1 ,1 06 and 3,385 samples, respectively. In both taxonomy datasets, there is a generally increasing trend in label count from coarser to finer ranks, accompanied by a generally decreasing trend in sample size. However, this pattern is not strictly monotonic across all levels.

**Fig 1.**
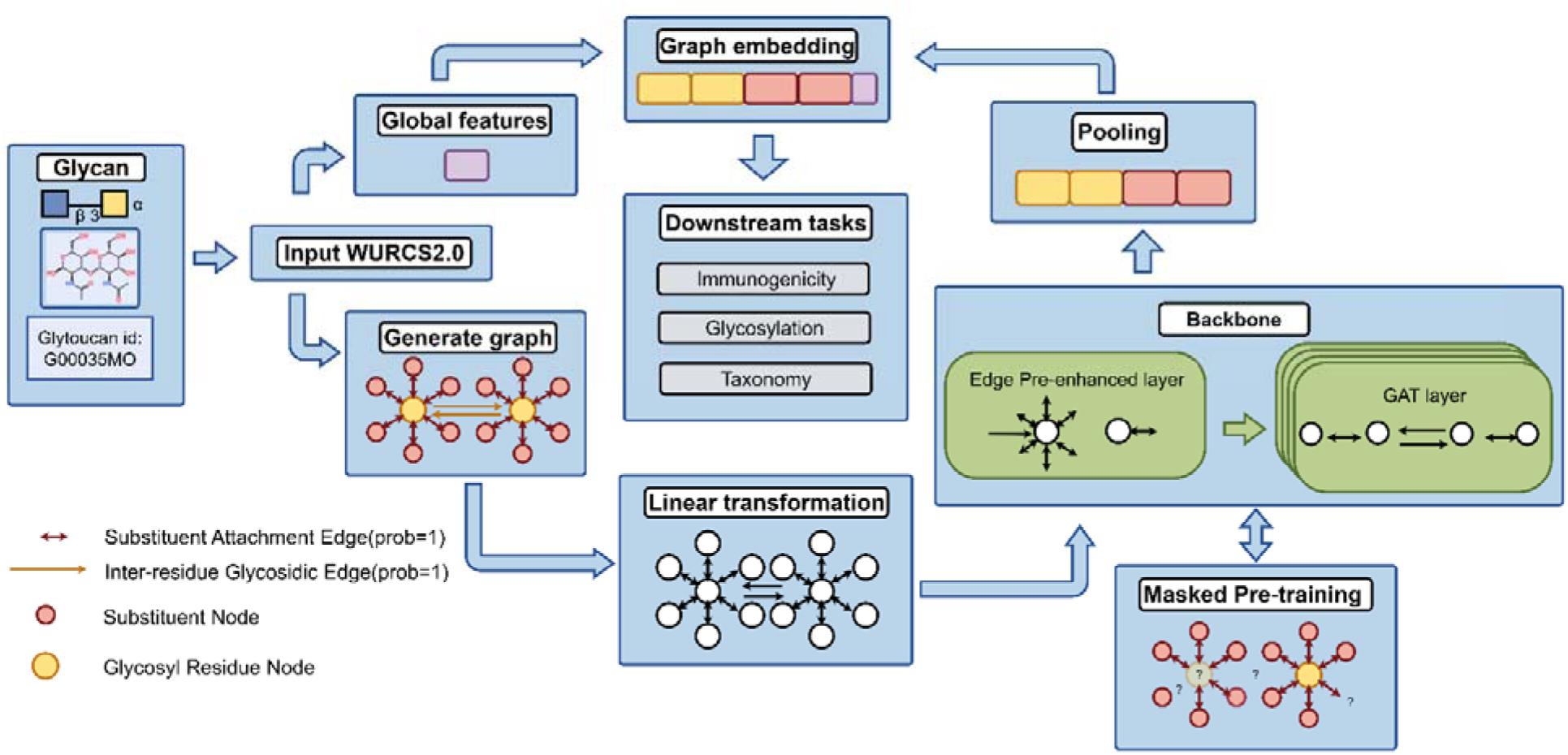
Schematic overview of the GlycanPUG framework. GlycanPUG – Probabilistic Uncertainty-aware Graph.

**Fig. 2.**
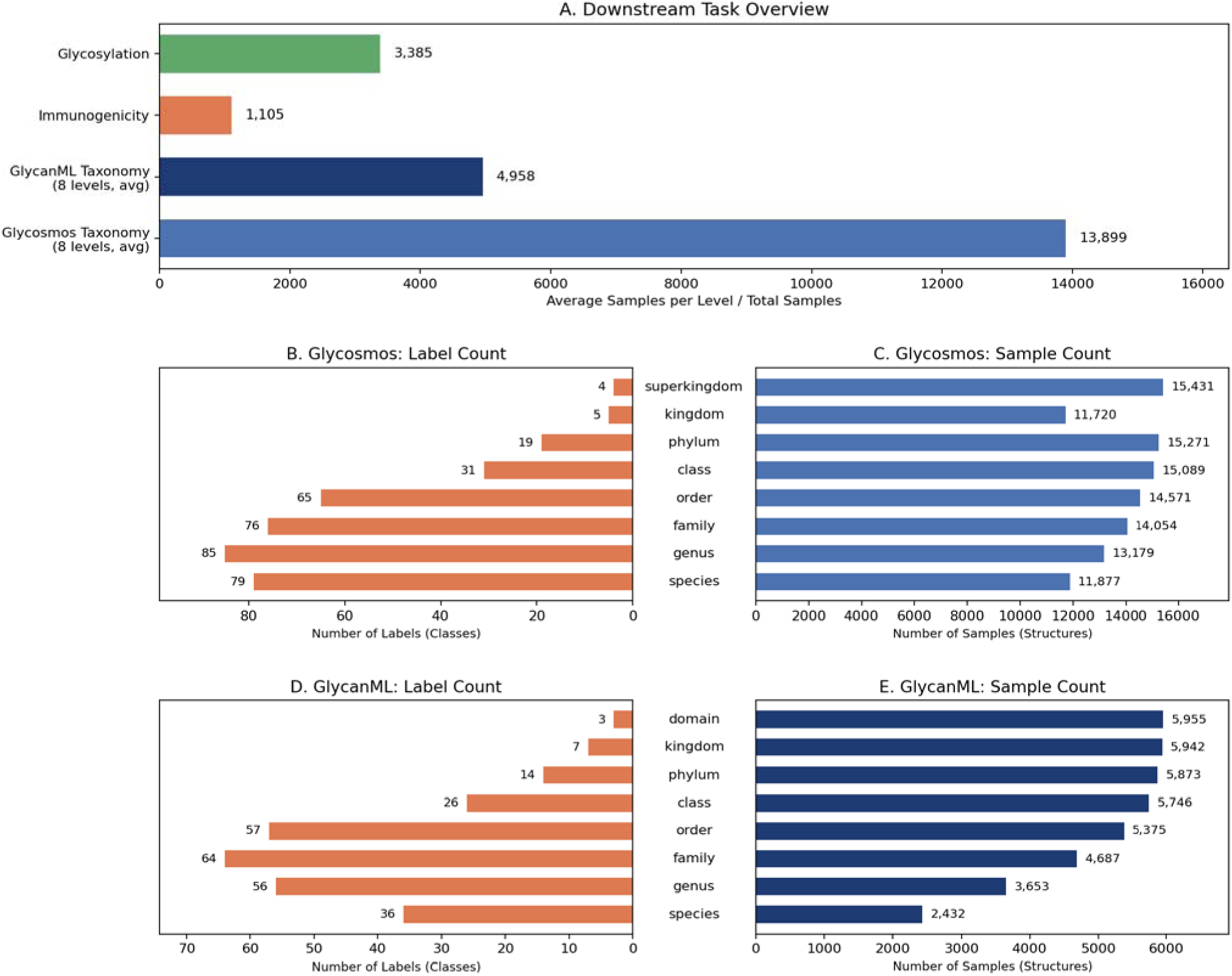
(A) Overview of four downstream tasks: glycosylation type classification, binary immunogenicity prediction, and multi-label taxonomy classification on GlyCosmos and GlycanML. For the taxonomy tasks spanning multiple hierarchical levels, average sample counts per level are reported. (B-C) GlyCosmos taxonomy dataset: label counts and sample counts across taxonomic ranks. (D-E) GlycanML taxonomy dataset: label counts and sample counts across eight taxonomic ranks.

This non-monotonicity is primarily driven by a frequency-based filtering step: labels occurring fewer than 30 times are pruned to ensure training reliability. At finer taxonomic ranks, although the theoretical number of distinct labels proliferates, a large proportion of these labels are rare and fall below the threshold, resulting in their removal.

### 3.2 Glycosylation Type Classification

Accurate identification of glycosylation types is fundamental to glycan functional annotation. Although GlycanML provides a standardized dataset for this task, the present study instead evaluates a GlyConnect-derived dataset containing structurally ambiguous entries. Therefore, the GlyConnect dataset(Alocci etal. 201 9), which retains structurally uncertain entries, was used after the preprocessing described in Section 2.2 to evaluate GlycanPUG. Glycan samples were retrieved from GlyConnect and categorized into three predominant classes: N-linked, O-linked, and free oligosaccharides.

### 3.3 Immunogenicity Prediction

To assess the baseline representational capacity of GlycanPUG on immunogenicity prediction, a head-to-head comparison was performed against GlycanGT on the binary immunogenicity prediction (IMM) task from the GlycanML benchmark. Macro AUPRC serves as the primary metric. All results are averaged over ten independent seeds (1 –1 0) to characterize variation across runs. As illustrated in Fig. 4(A), GlycanPUG achieves average performance that is on par with or slightly exceeds that of GlycanGT. The paired difference (GlycanPUG − GlycanGT) was computed, with a two-sided 95% confidence interval constructed accordingly, as detailed in Section 2. 7. The verdict for the IMM task was inconclusive (Fig. 4A,D), indicating that, for this task and these seeds, the data do not resolve a difference between the two models.

**Fig. 3.**
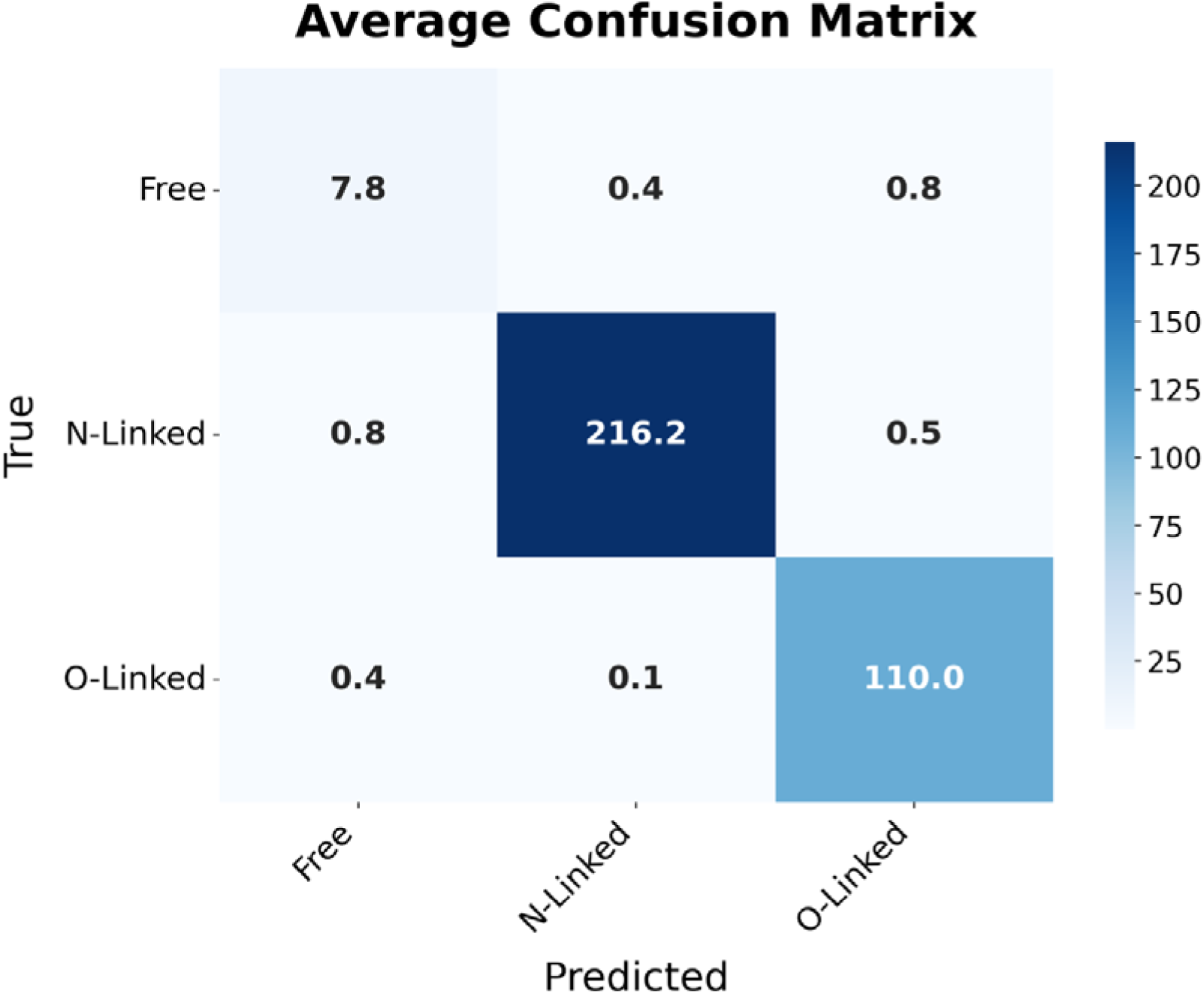
Evaluation of GlycanPUG on three-class glycosylation classification (N-linked, O-linked, free oligosaccharides) using the preprocessed GlyConnect dataset. Abbr: GlycanPUG – Probabilistic Uncertainty-aware Graph.

**Fig. 4.**
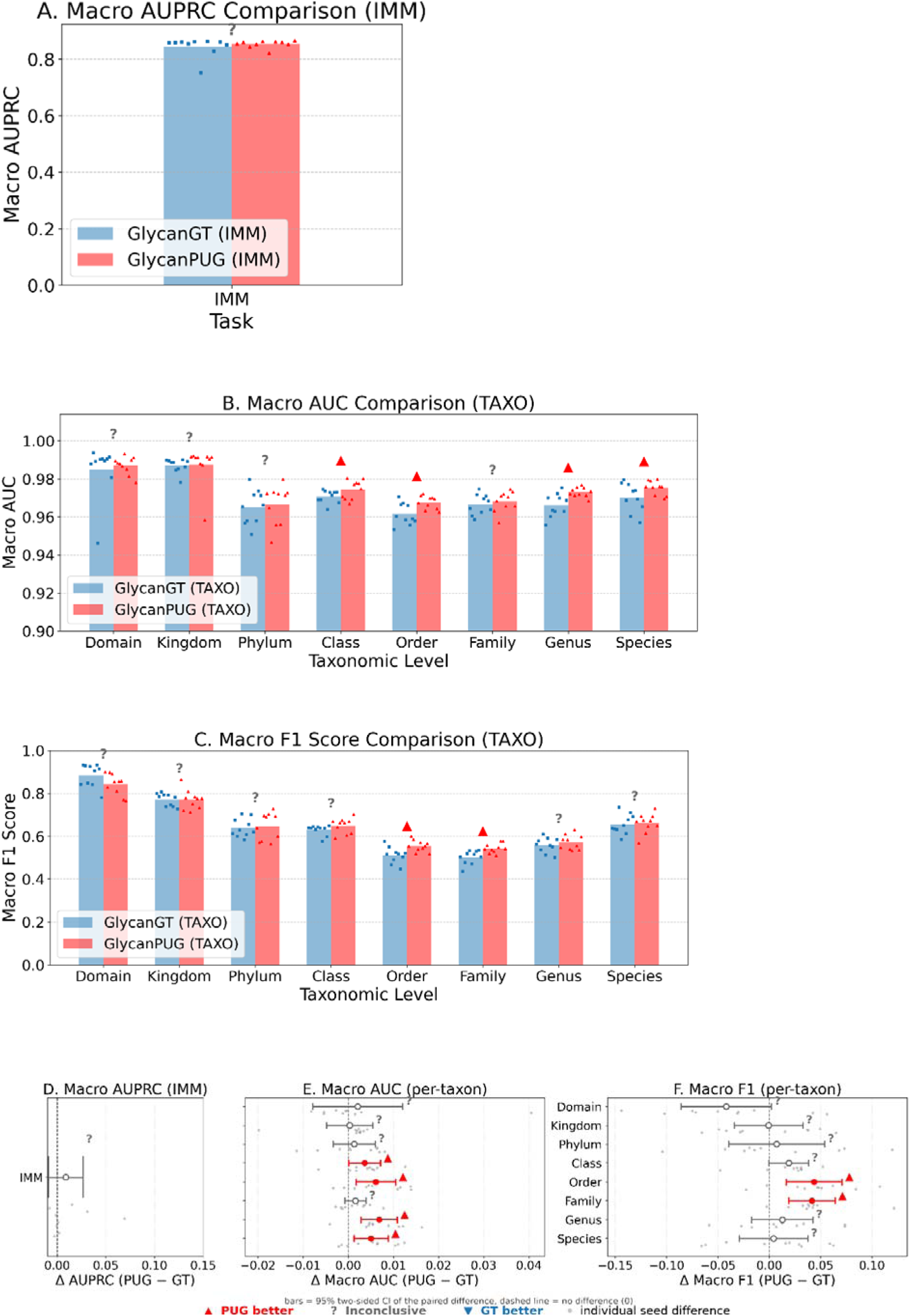
Benchmarking GlycanPUG against the GlycanGT (IUPAC-based) deterministic baseline on the GlycanML benchmark. Both models were used as fixed feature extractors with identical task-specific heads and evaluation splits, trained over ten independent seeds (1 –1 0). A Macro AUPRC for binary immunogenicity prediction (IMM); B, C Multi-label taxonomy classification across eight hierarchical ranks (Domain → Species): macro F1 (B) and macro AUC (C). D–F Paired differences (GlycanPUG − GlycanGT): AUPRC on IMM (D), per-rank macro AUC (E), per-rank macro F1 (F). Bars show two-sided 95% confidence intervals of the mean paired difference

### 3.4 Taxonomy Classification

To assess the baseline representational capacity of GlycanPUG on taxonomy classification, a head-to-head comparison against GlycanGT was performed. This evaluation covered multi-label taxonomy classification (TAXO) across all eight taxonomic levels (Domain to Species).

Performance is measured using Macro AUC and Macro F1 -score. All results are averaged over ten independent seeds (1 –1 0). As illustrated in Fig. 4(B; C), GlycanPUG achieves average performance that is on par with or slightly exceeds that of GlycanGT. To statistically validate this comparison, the paired difference (GlycanPUG − GlycanGT) was computed, with a two-sided 95% confidence interval constructed accordingly, as detailed in Section 2. 7. For Macro AUC (Fig. 4B,E), GlycanPUG was classified as “better” at the Class, Order, Genus, and Species levels, while results for Domain, Kingdom, Phylum, and Family were inconclusive. For Macro F1 (Fig. 4C, F), GlycanPUG was classified as “better” at the Order and Family levels, with the remaining levels inconclusive. No taxonomic level favored GlycanGT, supporting that GlycanPUG performs comparably to or better than the baseline across the full taxonomic hierarchy.

To further evaluate GlycanPUG on real-world structural uncertainty, multi-label taxonomy classification was performed on the GlyCosmos dataset. Unlike GlycanML, GlyCosmos entries inherently retain linkage and substituent uncertainties. Because no existing glycan-specific feature extraction model is capable of processing such inputs, this evaluation was conducted without a directly comparable external baseline. Results are presented in Fig. 5.

**Fig. 5.**
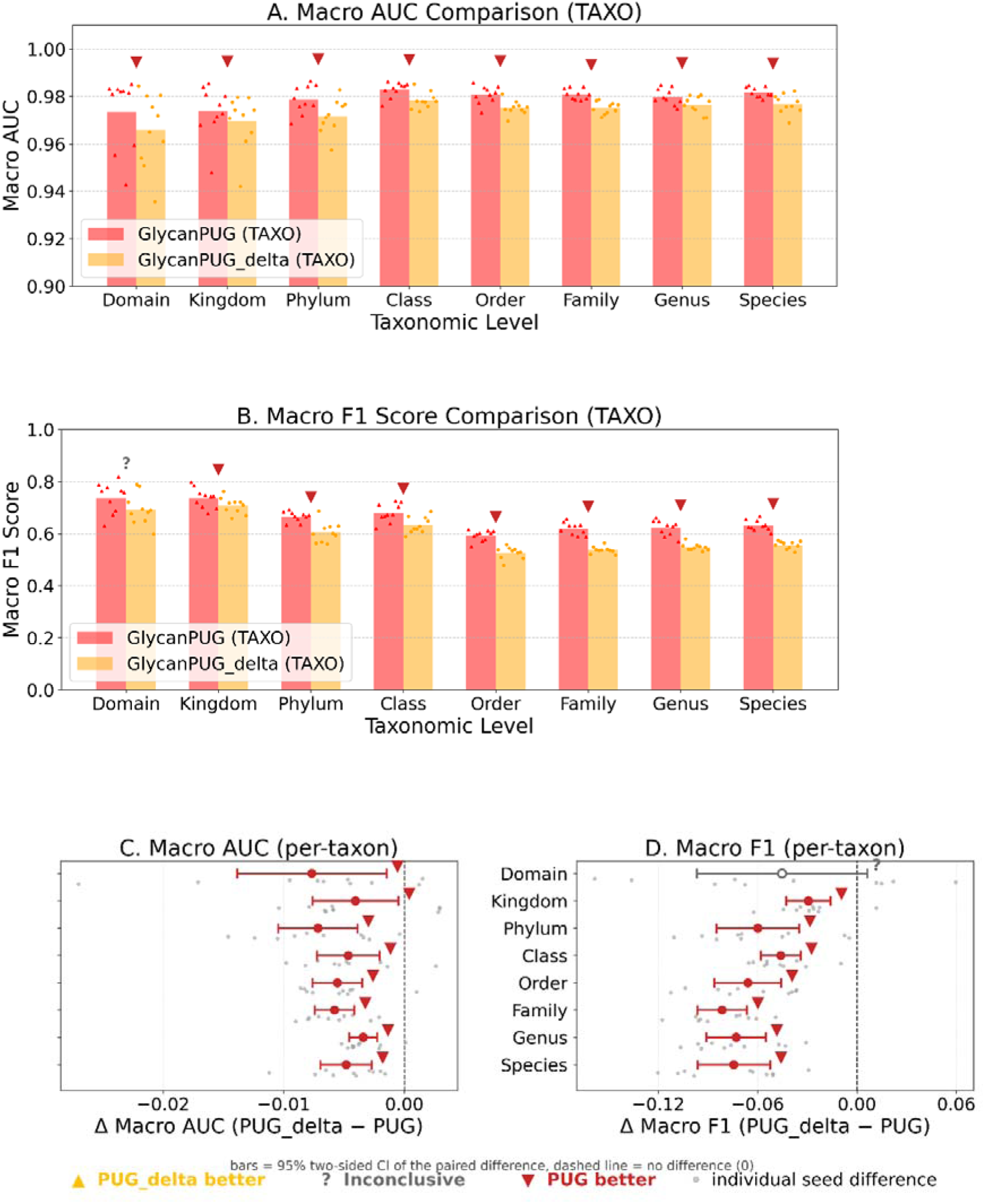
Ablation of the node pre-enhancement module: GlycanPUG vs. GlycanPUG_delta on multi-label taxonomy classification (GlyCosmos), across eight hierarchical ranks (Domain → Species), ten seeds (1 –1 0). A Macro F1 and B macro AUC per rank; Paired differences (GlycanPUG_delta − GlycanPUG) for macro AUC (C) and macro F1 (D).

**Fig. 6.**
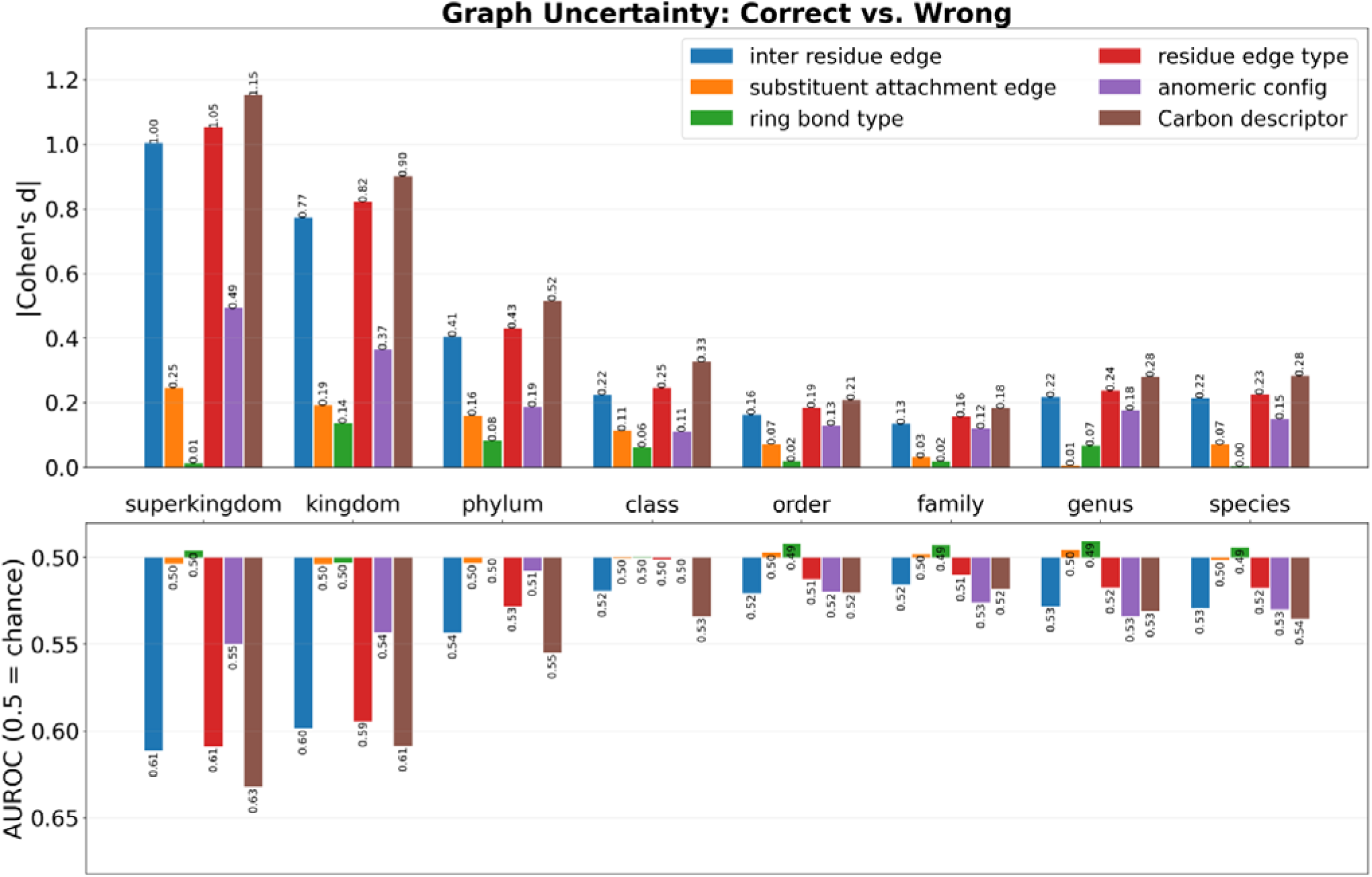
Upper panel: Absolute Cohen’s d for six uncertainty categories across eight taxonomic levels. Lower panel: Single-feature AUROC (0.5 = chance). Categories: inter-residue edge, substituent attachment edge, ring bond, residue edge type, anomeric, Carbon descriptor. Larger effect sizes (|Cohen’s d|) and discriminative power (AUROC) were observed at coarser taxonomic ranks for inter-residue edge, residue edge type, and Carbon descriptor, indicating associations with prediction errors at coarser taxonomic ranks. Ring bond type had small effect sizes across ranks; anomeric uncertainty had |d| = 0.55 at the coarsest rank. The results describe associations and do not establish the benefit of resolving uncertainty. Abbr: AUROC – Area Under the Receiver Operating Characteristic Curve.

### 3.5 Model Ablation

To quantify the contribution of the node pre-enhancement module, the full GlycanPUG model was compared against its ablated variant,. GlycanPUG_delta (which lacks the node pre-enhancement component). Multi-label taxonomy classification was performed on the GlyCosmos dataset across all eight taxonomic levels (Domain to Species). Performance was measured using Macro F1 -score and Macro AUC, with results averaged over ten independent seeds (1 –1 0).

As illustrated in Fig. 5, for Macro F1 -score, the comparison was inconclusive at the Domain level. At all other levels (Kingdom, Phylum, Class, Order, Family, Genus, and Species), GlycanPUG outperformed GlycanPUG_delta. Similarly, for Macro AUC, GlycanPUG consistently outperformed GlycanPUG_delta across all eight taxonomic levels. The per-taxon difference plots further confirm that the paired differences in both Macro AUC and Macro F1 are predominantly negative (PUG_delta − PUG < 0), with 95% two-sided confidence intervals largely favoring the full GlycanPUG model. Overall, these results confirm that the node pre-enhancement module contributes to predictive performance and that GlycanPUG consistently outperforms its ablated variant across most taxonomic ranks.

### 3.6 Association Between Structural Uncertainty and Prediction Errors

The relationship between graph-level uncertainty and classification errors was analyzed on the GlyCosmos taxonomy task (Fig. 6). Six uncertainty categories—inter-residue edge, substituent attachment edge, ring bond type, residue edge type, anomeric, and Carbon descriptor—were evaluated across all eight taxonomic levels.

Larger effect sizes (|Cohen’s d|) were observed at coarser taxonomic ranks for inter-residue edge, residue edge type, and Carbon descriptor. In the AUROC analysis (0.5 = chance), however, all six categories converged to near-chance values at finer taxonomic ranks, indicating limited marginal discrimination by these features at finer ranks in this analysis. Separately, ring bond type showed small effect sizes across ranks, whereas anomeric config had a larger effect size at the coarsest rank. These observational associations do not establish whether resolving a particular uncertainty would improve predictive performance.

## 4 Discussion

GlycanPUG is a pretrained graph model that represents supported glycan structural ambiguities. On the GlycanML benchmark (Xu, Geng *et al*. 2025), performance competitive with GlycanGT (Kitani *et al*. 2026) was achieved in multi-label taxonomy classification and binary immunogenicity prediction, with no statistically resolvable difference on the GlycanML benchmark. On GlyConnect (Alocci *et al*. 201 9) and GlyCosmos (Yamada *et al*. 2020), uncertain glycan entries were effectively handled by GlycanPUG. Ablation of the node pre-enhancement module caused a pronounced drop in macro F1 -score, supporting its contribution under the evaluated setting. These results support learning embeddings from glycans with supported structural ambiguities. The present experiments do not establish that the edge weights are calibrated probabilities or that the model performs inference over a joint distribution of valid glycan structures. Additionally, a post-hoc uncertainty–error correlation analysis revealed that the association of structural uncertainties with prediction errors varied substantially across uncertainty types.

Machine learning-driven glycan representation learning has developed substantially in recent years, progressing from early models like SweetTalk (Bojar 2020) for sequence-based language modeling and SweetNet (Burkholz, Quackenbush, and Bojar 2021 ) for graph convolutional networks, to more recent frameworks like GlycanAA (Xu, Song et al. 2025) for all-atom heterogeneous graph pretraining and GlycanGT for graph transformers. Among the latter, GlycanAA was developed for all-atom glycan modeling, employing heterogeneous graphs and hierarchical message passing to capture local-to-global dependencies, reporting strong performance on the GlycanML benchmark. Subsequently, GlycanGT introduced the graph transformer architecture with global self-attention to model long-range dependencies, with strong reported benchmark performance.

Despite the success of existing models, some preprocessing workflows restrict inputs to resolved structures. GlycanGT has made improvements by expanding enumerable uncertain linkages (e.g., “a1 -3/5”) into multiple graph instances, thereby increasing the number of pre-training samples. However, when extracting glycan features, it can still only process fully determined samples. Furthermore, enumeration generates alternative candidate structures; if their provenance and dependence are discarded, the representation may fail to distinguish a resolved linkage from one possible interpretation of an ambiguous linkage. Evaluated on 1 8,464 biologically derived glycans from the GlyCosmos database, the evaluated IUPAC-based workflow processes 7,952 entries (43. 1 % coverage; see Table 1 ), leaving many structurally ambiguous entries unprocessed.

**Table 1.**
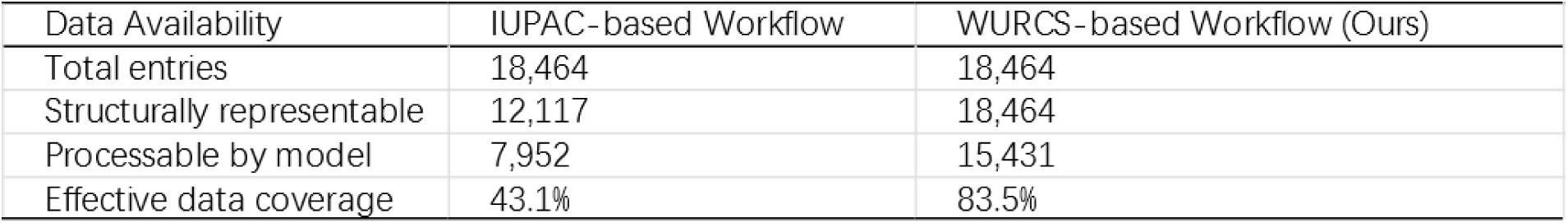
Data coverage on GlyCosmos-derived biologically sourced glycans (N=1 8,464). Comparison of IUPAC-based deterministic and WURCS-based workflows in total entries, structurally representable entries, model-processable entries, and effective coverage. Abbr: IUPAC – International Union of Pure and Applied Chemistry; WURCS – Web3 Unique Representation of Carbohydrate Structures.

By adopting WURCS 2. 0 (Matsubara *et al*. 201 7)—a canonical, database-centric notation now used by GlyTouCan (Tiemeyer *et al*. 201 7)—GlycanPUG broadens processable coverage in the evaluated cohort. The model uses weighted candidate edges and uncertainty attributes within a heterogeneous graph encoder (see Methods). Its intended contribution is the representation and use of supported structural ambiguities, rather than a new general-purpose graph attention operator. This design avoids explicit enumeration of complete candidate graphs and retains supported ambiguities within a single graph representation (effective data coverage: 83.5%; Table 1 ). Compositional decoupling was enabled by the primitive-level granularity of WURCS 2.0: each monosaccharide was decoupled into a “glycosyl residue node” and “substituent node”s, and chirality was explicitly represented as an edge attribute. This decomposition provides a feature vocabulary based on constituent structural attributes. The contribution of the node pre-enhancement module was evaluated through ablation: prior to each HeteroGNN layer, edge features were aggregated, weighted by their assigned values, and fused into the nodes, thereby isolating edge-to-node information transfer from joint message passing. Removal of this module led to significantly degraded performance—GlycanPUG outperformed GlycanPUG_delta across all eight taxonomic levels for Macro AUC, and at seven of eight levels for Macro F1 -score (inconclusive only at the Domain level), with two-sided 95% confidence intervals consistently favoring the full model. Additionally, post-hoc analysis revealed that associations between structural uncertainty and prediction errors varied by taxonomic rank. At coarser taxonomic ranks, inter-residue edge, residue edge type, and Carbon descriptor uncertainties were strongly associated with misclassification, whereas at finer ranks (Genus–Species) all six categories converged to near-chance discrimination, indicating limited marginal discrimination by the six aggregate uncertainty features at these ranks. Ring bond type showed small effect sizes across all ranks. These associations do not quantify the causal benefit of resolving structural uncertainty.

This framework is primarily optimized for linear and branched oligosaccharides; variable-repeat polysaccharides (∼4% of GlyTouCan) feature an indeterminate polymerization degree n, posing a fundamental limitation for canonical graph construction, and thus were omitted from the dataset, rendering the framework incapable of modeling polysaccharides with variable repeat units. Quantitative binding prediction is challenged by dataset heterogeneity: while internal batch normalization is performed by some workflows to retain more samples, variations in key covariates (e.g., spacer length, assay concentration, glycan types) are often overlooked, risking that experimental artifacts are confounded with genuine signals; consequently, direct prediction of absolute binding affinities is not performed. From a modeling perspective, probabilistic graphical frameworks are theoretically capable of capturing repeatingÖunit distributions but are hindered by data scarcity and computational costs; alternative hierarchical encoding schemes are similarly constrained by supervised data scale. Therefore, such methods are only conceptually discussed in this study, with specific implementations deferred to future work.

## Computational Environment

All experiments were conducted on a workstation running Windows 11 equipped with a single NVIDIA GeForce RTX 5060 GPU (CUDA 1 2.8). The software environment was based on Python 3.1 3.9, with PyTorch 2.9.0 (CUDA 1 2.8 build) as the deep learning framework and PyTorch Geometric 2.7.0 for graph neural network implementation. Data processing and evaluation relied on NumPy 2.3.4, pandas 2.3.3, SciPy 1 .1 6.3, and scikit-learn 1 .7.2, while multi-label stratified cross-validation was performed using iterative-stratification 0.1 .9. All figures were generated with Matplotlib 3.1 0.7 and Seaborn 0. 1 3.2. Codes and model weights uploaded on (https://github.com/wangyefei958dot/GlycanPUG.git.)

## Author contributions

Yefei Wang (Conceptualization [Lead], Data curation [Lead], Formal analysis [Lead], Investigation [Lead], Methodology [Lead], Software [Lead], Validation [Lead], Visualization [Lead], Writing—original draft [Lead], Writing—review & editing [Lead]).

## Acknowledgements

The authors thank the National Natural Science Foundation of China (NSFC) (Nos. 821 30099 and 22477030), Science and Technology Commission of Shanghai Municipality (grant No.24DX1 400200), the International Cooperation Program of Shanghai Science and Technology (No.23490711 600), the Fundamental Research Funds for the Central Universities (222201 71 7003), the Programme of Introducing Talents of Discipline to Universities (B1 601 7), the National Natural Science Foundation of Shanghai Science and Technology (No.24ZR1 41 5400), the Shanghai Oriental Talents youth Program (No. QNKJ202401 0), Fundamental and Interdisciplinary Disciplines Breakthrough Plan of the Ministry of Education of China (JYB2025XDXM404), the Open Funding Project of the State Key Laboratory of Fine Chemicals, Dalian University of Technology (KF 2402 and KF 2507), State Key Laboratory of Chemo/Biosensing and Chemometrics, Hunan University, Changsha 41 0082, P. R. China, the State Key Laboratory of Medicinal Chemical Biology (2026005), NanKai University for financial support. The Research Center of Analysis and Test of East China University of Science and Technology was gratefully acknowledged for assistance in analytical experiments.

## Abbreviations

AUC: Area Under the Curve
AUROC: Area Under the Receiver Operating Characteristic Curve
AUPRC: Area Under the Precision-Recall Curve
EMA: Exponential Moving Average
F1: F1 -score (harmonic mean of precision and recall)
CI: Confidence Interval
GAT: Graph Attention Network
GCN: Graph Convolutional Network
GNN: Graph Neural Network
GlycanPUG: Probabilistic Uncertainty-aware Graph
GlycanPUG_delta: Probabilistic Uncertainty-aware Graph (ablation)
HGNN: Heterogeneous Graph Neural Network
IUPAC: International Union of Pure and Applied Chemistry
MLM: Masked Language Modeling
ROC: Receiver Operating Characteristic
WURCS: Web3 Unique Representation of Carbohydrate Structures
TAXO: Taxonomy

## Notes

https://github.com/wangyefei958-dot/GlycanPUG.git

### Competing Interest Statement

The authors have declared no competing interest.

