## Supplementary Material for "GlycanPUG: WURCS-Based Heterogeneous Graph Pretraining for Structurally Ambiguous Glycans"

**A. Preliminaries for WURCS 2.0**

Although WURCS 2.0 ^1^ is an established standard, the model architecture is directly derived from its syntax. Therefore, a clear parsing explanation is essential for understanding the graph construction, and is provided below to describe the model inputs rather than replace the official WURCS 2.0 specification.

GlyTouCan ID: G00035MO

"WURCS=2.0/2,2,1/[a2112h-1a_1-5_2*NCC/3=O][a2122h-1b_1-5_2*NCC/3=O]/1-2/a3-b1"

Parse the top-level WURCS sections using their syntax; slashes within MAP codes must not be treated as top-level delimiters.

WURCS= Version/ Unit Count/ UniqueRES List/ RES Sequence/ LIN List.

**A.1 Version and Unit Count**

The prefix 2.0 denotes the WURCS version. The subsequent triplet 2,2,1 constitutes the Unit Count area, encoding three foundational parameters: UniqueRES count (number of unique residue types), RES count (total number of residues), and LIN count (the number of LIN declarations).

**A.2 UniqueRES area**

The bracketed segments define the residue templates. Taking the first template [a2112h-1a_1-5_2*NCC/3=O] as an example, it can be decomposed into four components:

Skeleton Code (a2112h): The six-character code denotes a six-carbon monosaccharide residue. For convenience, these characters are provisionally referred to as the Carbon descriptor feature, which encodes the local connectivity and stereochemical information for each carbon;

Anomeric configuration (1a): The number identifies carbon 1 as the anomeric position, and a denotes the α configuration.

Ring formation (1-5): The segment defines the ring-forming carbon atoms within this individual residue (using local backbone carbon numbering); since the ring type is not explicitly stated, the omitted MAP represents an ether linkage under the WURCS omission rules.

Substituent (2*NCC/3=O): The suffix signifies that an NCC/3=O group is attached at position 2.

**A.3 RES Sequence and LIN List**

The RES Sequence section 1-2 indicates the presence of two residues, mapping Residue 1 to Template 1 and Residue 2 to Template 2. The LIN List section a3-b1 then specifies the connectivity: a (Residue 1) links to b (Residue 2) via position 3 of Residue 1 and position 1 of Residue 2.

**A.4 Handling uncertainty**

Within the template area, WURCS employs specific placeholder symbols to denote incomplete structural information. The character x is used to indicate unknown stereochemistry—appearing in both the Carbon descriptor code (e.g., replacing a specific carbon-bonding descriptor) and the anomeric configuration prefix (e.g., x instead of a or b). Conversely, the symbol “?” denotes an unknown carbon position, such as an undetermined ring-forming locus or substituent attachment site. These placeholders enable WURCS to represent partially characterized glycans while preserving a valid structural framework.

In the linkage area, “?” similarly signifies an unknown connection site on a residue, while the pipe | expresses a disjunction over linkage targets. For example, the notation “a?-b?|c?}” indicates that Residue 1 is linked to either Residue 2 or Residue 3.

In rare cases, the linkage region may contain uncertain notations such as “a?|b?|c?}*OCC/3=O”, indicating the presence of an OCC/3=O group attached to one of residues 1, 2, or 3, with an unspecified attachment position. While such entities are counted in the standard Lin count, true linkage count is introduced to exclude these uncertain substituents, thereby quantifying inter-residue linkage declarations, including declarations with unresolved positions or connectivity.

**B. Conversion from IUPAC to WURCS**

Since the GlycanML ^2^ dataset only provides IUPAC notations, the glycan structures were converted to WURCS 2.0 format using GlycanFormatConverter ^3^ via the GlyCosmos ^4^ API. Out of 13,209 total samples, 6,608 were successfully converted in the initial round. Terminal carboxylated IUPAC residues such as GlcA were found to go unrecognized by the converter. To address this, "?F" was adopted as a placeholder—fluorine substituents being extremely rare in biological contexts, and uncertainly positioned fluorine substitutions rarer still—and GlcA was correspondingly rendered as an intermediate form, e.g., "Glc?F". Patterns matching "?F" were then identified and corrected to their proper forms. Substituents such as OMe, OP, and OS without specified linkage positions were uniformly replaced with "?Me", "?P", and "?S", respectively. However, for OAc, the converter could not interpret “?Ac”, so “?F” had to be retained as a placeholder. NeuNAc was substituted with the converter-compatible term Neu5Ac. GlyCosmos also offers an API to convert WURCS 2.0 back into IUPAC notation, which was used for a round-trip consistency check. Nevertheless, because IUPAC itself does not prescribe a fixed main chain, the regenerated IUPAC strings often differed from the originals in terms of branching order and side-chain arrangement. Therefore, a naïve character-count comparison (brackets ignored) was insufficient to establish structural equivalence. Instead, round-trip fidelity was verified by parsing both the original and the regenerated IUPAC strings into residue trees and comparing the resulting structures**,** where nodes encode monosaccharide identities together with their ring substituents and edges encode anomeric configuration and linkage positions. A canonical signature is computed for each tree by sorting child-node signatures, so that structurally identical glycans are recognized as equivalent regardless of branching order or side-chain arrangement. An input–output pair was accepted as consistent only when their root signatures matched; otherwise, residue-level and linkage-level differences were reported.

Of 13,209 glycans in the GlycanML classification dataset, 9,387 (71.1%) were successfully converted and passed structural verification; of 1,320 in the GlycanML immunogenicity dataset, 1,105 (83.7%) were accepted. The remaining failures were analyzed and categorized (Table S.1). The dominant cause was residues or substituents not covered by the GlycanFormatConverter vocabulary (50.5% in classification; 25.6% in immunogenicity, including HTTP 500 errors which were confirmed to originate from unrecognized input). Other causes included placeholder collisions (3.3%), multiple modifications on a single carbon (1.8%), and ketose anomeric ambiguity (1.6%). The tree-isomorphism verification rejected only 1.73% (classification) and 0.72% (immunogenicity) of round-trip conversions, confirming that the accepted samples represent genuine structural equivalences. Labels with ≥30 occurrences were retained for the multilabel classification task.

**C. Graph Construction**

The following sections describe the features of each node and edge type.

**C.1 Carbon Count, Anomeric Position, and Anomeric Configuration**

These three features — carbon count, anomeric position, and anomeric config — are straightforward to interpret. However, several edge cases warrant clarification.

Unknown anomeric position is encoded by the "?" character in WURCS 2.0 and has been added as a dedicated token to the model vocabulary. Unknown anomeric config is encoded by the "x" character, which has also been added as a dedicated token.

Absence of anomeric information is handled differently depending on residue type. For known residues, the omission of anomeric descriptors in WURCS 2.0 indicates that no anomeric center exists (e.g., open-chain alditols); both features are assigned the "N/A" token. For fully unspecified residues denoted by the Q placeholder, two cases arise: [<Q>] (no anomeric suffix) means the structural details — including whether an anomeric center is present — are entirely unknown, and both features receive the "unknown" token; [<Q>-<pos><config>] (with anomeric suffix) partially specifies the anomeric center despite residue identity being unknown, where the suffix follows the standard WURCS anomeric descriptor syntax (a positional digit or ? followed by a config character such as a, b, x, etc.) and is parsed directly into the corresponding position and config tokens.

**C.2 Ring Bond Type and Double Bonds**

Ring bond type and double bonds are attributes of glycosyl residue nodes. Before each GAT layer, these features are concatenated with the original Glycosyl Residue Node embeddings. Since a single residue may contain multiple rings or double bonds, their representations are aggregated as follows: For ring bond type, entries are formatted as "start pos-end pos" based on template positions (e.g., retaining uncertain positions as "1-?"). For double bonds, entries use the format "pos=pos", based on template positions. In cases of multiple instances, the corresponding embeddings are averaged via arithmetic mean. For residues lacking ring structures, ring bond type is set to "N/A"; similarly, double bonds is set to "none" if absent. When structural information is entirely unavailable, both attributes default to "unknown".

**C.3 Residue edge type and edge prob (for** **Inter-residue Glycosidic Edge)**

In message-passing GNNs, reverse edges are often generated by simply transposing the edge index matrix. However, this approach assumes edge attributes are either symmetric or irrelevant to direction. In the present setting, Residue edge type is inherently directional—the reverse of a "3-1" linkage is semantically a "1-3" linkage, not a duplicate "3-1". Because standard index reversal leaves edge-type features untransformed, reverse edges were explicitly constructed with their correct type labels, thereby preserving accurate directional information for the model.

Consider the LIN list fragment a2|b?}-c1, which denotes that residue c at position 1 is connected to either site-2 of residue a or an undetermined site of residue b. The parser expands the disjunction | into two separate edges while preserving the positional uncertainty “?” as an edge attribute, specifically generating a forward edge from a to c tagged (2-1) with an assigned weight of 1.0, and another from b to c tagged (?-1) with an assigned weight of 0.5. The corresponding reverse edges are similarly produced: c to a tagged (1-2) with weight 0.5, and c to b tagged (1-?) with weight 0.5. Only the alternative-target relationship introduced by | triggers edge expansion in this implementation. In contrast, fully expanding the “?” uncertainty (e.g., enumerating every candidate site on residue b) would fragment the assigned weight across many candidates, potentially attenuating individual edge contributions. By preserving “?” as a symbolic tag rather than expanding it, the representation avoids explicit enumeration of every candidate carbon position.

**C.4 substituent, Carbon descriptor, edge prob (for substituent Attachment Edge) and carbon position type**

All carbon positions are initialized with a default oxygen substituent "O". Accordingly, substituent nodes representing "O" are created based on the number of carbons and connected to the parent residue nodes via an equal number of substituent attachment edges. When all connectivity is confirmed, the number of such substituent nodes and their associated substituent attachment edges equals the carbon count of the residue. These edges carry three features: carbon position type (indicating the index of the connected carbon within its residue), Carbon descriptor (the character at the corresponding position in the residue's skeleton code, encoding local connectivity and stereochemistry at that carbon), and edge probability (initialized to 1.0 and later adjusted per competing uncertain substituents). This default "O" assignment is overridden either by special substituents inferred from stereo characters—for example, 'd'/'m' denote deoxygenation (H), while double-bond characters ('E/e/Z/z/F/f/n/N') map to "none"—or by explicitly declared substituents in the template (e.g., "2*OSO3").

A carbon position is considered available if it is not involved in ring closure, not engaged in a confirmed inter-residue glycosidic linkage, and not occupied by a confirmed non-O substituent; positions with uncertain connectivity (unknown linkage presence rather than uncertain linkage mode) also remain available. For position-uncertain substituents (including those in residue templates and linkage regions), their probability mass (encoded as the edge probability feature, weight = 1/N, where N is the number of available sites) is uniformly distributed across all available carbon sites; the allocated probability mass is deducted from the corresponding default O-substituent edges. If the total deducted amount at a given site reaches 1.0, the O-edge is fully consumed and removed—it never becomes negative. Multiple uncertain substituents may independently target the same carbon atom; Such competing edges were deliberately retained as an intentional approximation.

**C.5 Special notes**

Most ring closures occur via oxygen atoms, though non-oxygen atom-mediated ring formation is also observed in certain cases. In [a2122A-1b_1-5_2-3OC^XO/3CO/6=O/3C], the ring closure between positions 2 and 3 is formed through the OC^XO/3CO/6=O/3C moiety. This 2-3 linkage was designated as the ring closure information, while simultaneously treating both positions 2 and 3 as bearing the substituent OC^XO/3CO/6=O/3C. This representation aligns with the treatment applied to standard ring closures. For a typical 1-5 oxygen ring, independent O-substituent nodes are placed at both C1 and C5, with the 1-5 connection recorded as Ring Bond Type. Analogously, for a bridging substituent such as OC^XO/3CO/6=O/3C linking positions 2 and 3, separate nodes of that substituent are instantiated at each connected carbon, and the 2-3 linkage is designated as Ring Bond Type.

**C.6 Graph Representation and Input Pre-processing**

The elements of the graph, their associated features, pre-processing steps prior to GAT input, and corresponding embedding dimensions are summarized in Figure S.1.

**D. Pretraining Tasks**

The pretraining phase employs masked attribute prediction and edge-reconstruction tasks to learn glycan structural representations. To balance readability and completeness, the overall hyperparameters—including mask ratios, loss weights, loss functions, and standard masking/masking-phase configurations for all tasks—are systematically summarized in Table S.2. Meanwhile, the following paragraphs elaborate on the specific design rationales and non-trivial implementation details of several complex tasks.

In the carbon number prediction task, The carbon count type was masked and all substituent attachment edges were removed to eliminate trivial information-leakage pathways (e.g., recovery of the carbon count from the number of substituents or edges). However, the residue-level features—ring bond type and double bonds—were deliberately retained unmasked. Although these features exhibit a weak correlation with carbon count—for instance, a 1–5 ring closure indicates that the residue possesses at least five backbone carbons and is statistically more likely to be a six-carbon sugar—this relationship is not deterministic. Not all six-carbon sugars adopt a 1–5 ring (e.g., furanoses use a 1–4 ring), and a 1–5 ring can also appear in non-six-carbon residues (e.g., modified heptoses). Similarly, double-bond positions indicate unsaturation but do not encode the total carbon count. Consequently, these features only provide ambiguous lower-bound cues rather than an exact carbon count. Masking them would discard chemically meaningful structural context that aids the encoder in learning general glycan representations, while yielding negligible benefit in preventing shortcut learning. They were therefore deemed acceptable as inputs to the carbon-number prediction task.

In theory, the double bonds prediction task should adopt multi-label classification, consistent with the ring bond type prediction task. However, after filtering the entire pre-training dataset, only the bond type "4=5" is present. Consequently, the task effectively reduces to a binary classification problem distinguishing between "none" and "4=5".

The edge existence pretraining task follows the principle of existence being independent of edge attributes: it predicts whether a connection exists between two residues rather than its specific linkage type. Question-mark annotations reflect only positional uncertainty; therefore, edges with probability exactly 1.0 and a single candidate are still treated as deterministic positive samples. Molecular graphs with fewer than 4 residues are excluded. Within a batch, if no eligible deterministic edge exists, or if Bernoulli sampling at p=0.15 per candidate edge yields zero hits, the task is skipped for that batch. Positive samples are deterministic edges with probability exactly 1.0, including single-candidate question-mark edges; both the masked edge and its reverse edge are simultaneously removed from the message-passing graph, with only the forward direction counted as positive. For each masked positive edge, at most one negative edge is generated per endpoint, yielding a normal negative-to-positive ratio of 2:1. During generation, with 70% probability the sampler attempts to select a hard negative from the opposite endpoint's neighbors, falling back to uniform random within the same graph if that branch yields no valid candidate; the remaining 30% of cases directly use uniform random within the same graph. All branches share unified constraints: the candidate pair must not be an existing edge in the graph—this set includes edges with probability below 1.0 and their reverse edges, ensuring negatives are definitively non-existent connections. When an endpoint has no candidate through either the hard-negative path or the random-fallback path, that endpoint produces zero negatives and the positive edge degrades to a 1:1 ratio; when both endpoints yield no candidates, the positive edge is excluded from the loss to prevent positive-only bias. Although this theoretically wastes one training signal, such cases were not observed in the actual training data. Negative samples are restricted to within the same molecular graph; intra-batch deduplication via a used-pair set ensures zero duplicates, scoped to a single forward-pass batch. Equally-weighted positive and negative samples are concatenated and optimized with binary cross-entropy; negatives account for approximately two-thirds of the total loss, with minor per-batch fluctuations due to degeneracy handling.

**E. Uncertainty Feature Definitions**

The following six aggregate metrics quantify structural ambiguity at the graph level, extracted independently for each test sample. Each metric produces a scalar in [0,1), where higher values indicate greater uncertainty.

**E.1 Inter-residue edge uncertainty**

Measures the proportion of unconfirmed residue-to-residue connections. Computed as
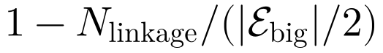
, where
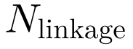
 is the number of confirmed linkages (parser metadata true linkage count) and
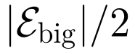
 is the number of undirected inter-residue edges. Since each inter-residue edge is explicitly added together with its reverse edge, the raw edge count is twice the number of underlying bonds; Raw edge counts were therefore divided by 2, thereby ensuring that the denominator captured only unique residue–residue connections. A value of 0 means all connections are resolved; values near 1 indicate high ambiguity in residue connectivity.

**E.2 Substituent attachment edge uncertainty**

Measures ambiguity in substituent placement. Computed as
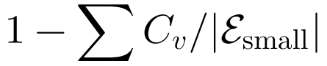
, where
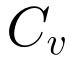
 is the confirmed carbon-position count of residue
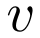
 and
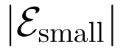
 is the total number of Substituent Attachment Edge (including doubt-generated candidates). Higher values indicate more unresolved attachment sites.

**E.3 Ring bond type uncertainty**

Proportion of ring-bond descriptors containing an unresolved placeholder “?”. Computed as the fraction of ring-bond entries where ring bond type contains “?”.

**E.4 Residue edge type uncertainty**

Mean score of unresolved placeholders in the connection-type descriptor of undirected inter-residue edges. Each edge receives
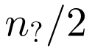
 points, where
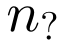
 is the count of “?” characters in the edge-type string (one “?” → 0.5, two “?” → 1.0). The mean across all undirected edges is reported. Each descriptor has two slots (donor and acceptor), and because the forward and reverse edges duplicate the same two slots, both directions contribute identically, so averaging over directed edges yields the same proportion as over unique bonds.

**E.5 Anomeric configuration uncertainty**

Proportion of residue-level nodes whose anomeric configuration is marked as unresolved (x).

**E.6 Carbon descriptor uncertainty**

Proportion of Carbon descriptors marked as unresolved (x), computed as a probability-weighted ratio. Each Substituent Attachment Edge contributes its assigned probability
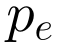
 to the denominator; edges with stereo type = "x" contribute
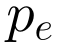
 to the numerator. This weighting ensures that doubt candidate edges (with
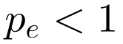
) are counted proportionally. The score is therefore equivalent to the fraction of probability mass landing on x slots in the carbon descriptor.

**E.7 Association analysis**

For each metric and taxonomic rank, correctly classified and misclassified samples were pooled across ten random seeds (1-10), and distributions were compared using a one-sided Mann–Whitney U test (direction-aligned: misclassified > correct for all doubt-type metrics), Cohen's d (direction-aligned: misclassified − correct), and single-feature AUROC (treating the metric as a classifier for misclassification). Reported values are single estimates from the pooled sample, with mean ± standard deviation denoting within-group feature values (correct vs. misclassified), not across-seed variability.

**F Step-wise Sample Counts in Pretraining Data Preprocessing**

The preprocessing pipeline consists of a sequence of filtering and conversion steps applied to both the pretraining and downstream corpora. For the pretraining data, samples overlapping with downstream tasks were first removed by exact matching on canonical WURCS 2.0 strings to prevent data leakage (Step 1; 21,273 removed, 232,323 remaining). Glycans containing repeating structural units were then excluded, as their copy numbers are indeterminate and their heterogeneity is high (Step 2; 7,000 removed, 225,323 remaining). Next, a combined filtering step targeted two criteria: entries with uncertain notation in the Unit Count region—which records the essential compositional unit information of a glycan and whose ambiguity prevents reliable graph initialisation and feature counting (7,015 removed)—and samples carrying rare linkages (768 removed), for a joint reduction of 7,783 (Steps 3–4; 217,540 remaining). Within the rare-linkage category, nitrogen-mediated linkages are the most prevalent; they give rise to atypical local topologies in which three residues converge on a single nitrogen atom, which would require a dedicated nitrogen node type and bespoke edge definitions in the graph neural network, yet their abundance is too low to avoid severe class imbalance. Samples with excessive uncertain connectivity were then discarded by computing the ratio of the confirmed linkage count (Appendix A.4) to the total number of unique inter-residue glycosidic edges (Appendix C; directed edge count halved to cancel the explicit reverse edges) and removing any sample below 0.1 (Step 5; 21,763 removed, 195,777 remaining). Finally, an iterative low-frequency token filter was applied across multiple feature categories (Appendix C): any token occurring in fewer than 0.01% of entries was dropped, and the pass was repeated until no rare features remained, thereby tightening the retained vocabulary (Step 6; 28,757 removed). After all filtering steps, the initial 253,596 pretraining samples were reduced to 167,020; per-step counts are reported in the table S.3

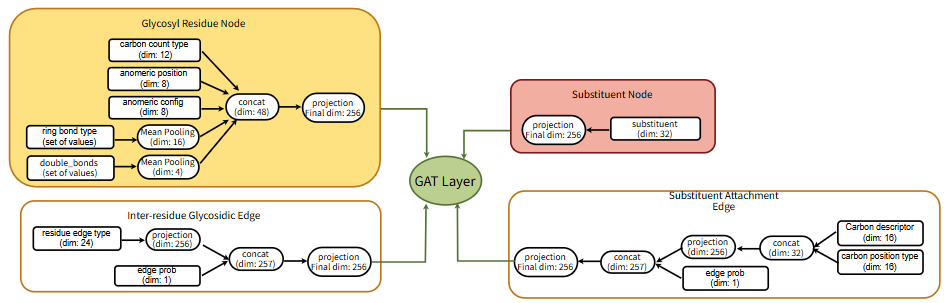

Fig S.1 Overview of graph elements, feature engineering, pre-processing steps, and embedding dimensions for GAT input.

Table S.1. Failure modes in two-round IUPAC->WURCS conversion

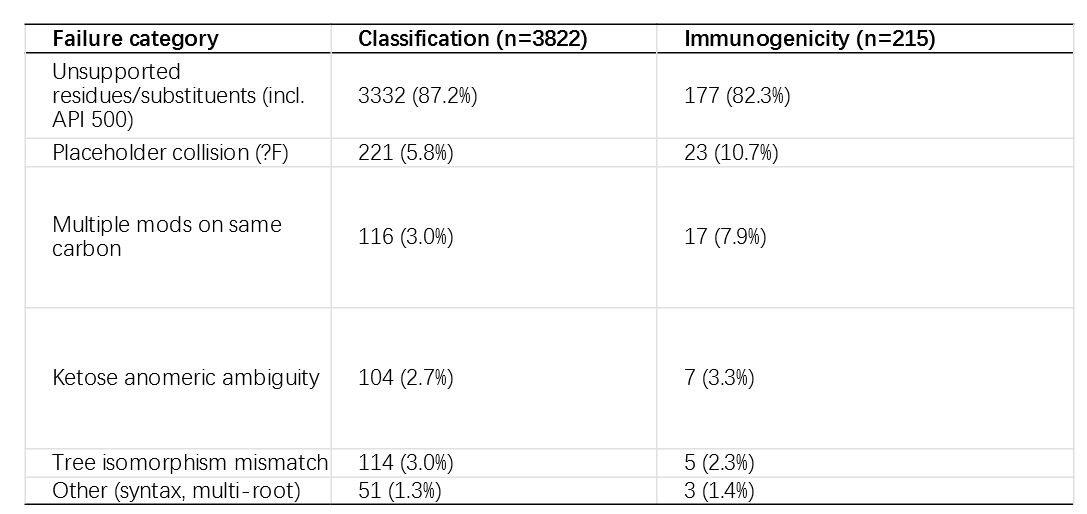

Table S.2 Hyperparameter configurations—including mask ratios, loss weights, loss functions, and special masking/training-phase settings—for all pretraining tasks, with the design rationales for complex tasks elaborated in Appendix D.

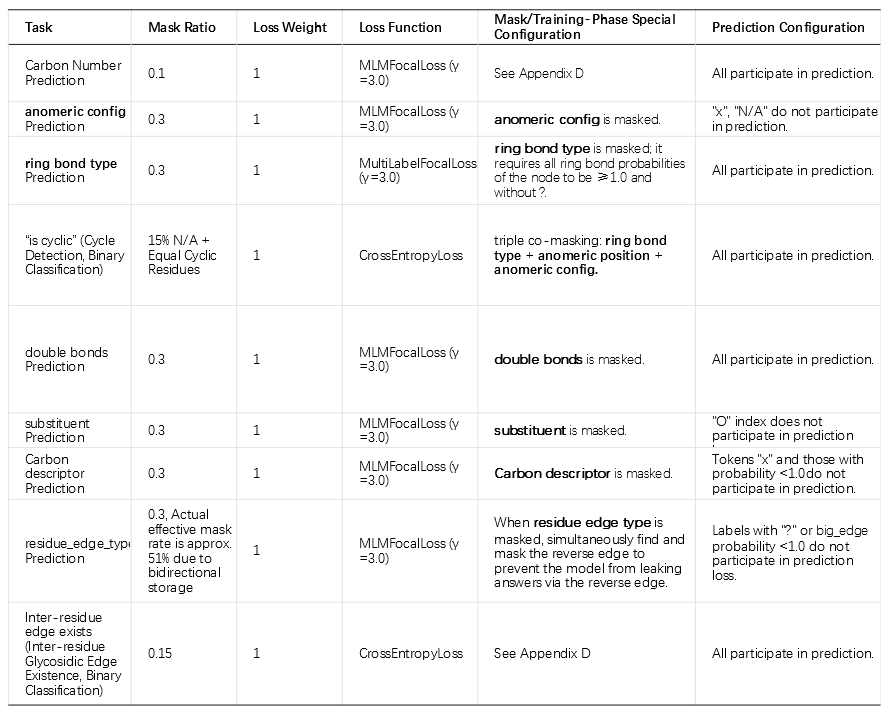

Table S.3 Preprocessing filtering steps and data counts

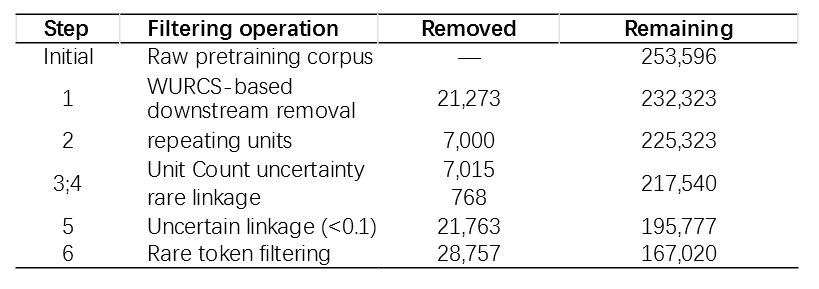
